# Inferring viral occurrence patterns through a synthetic data simulation

**DOI:** 10.1101/2021.07.13.452220

**Authors:** Ville N. Pimenoff, Ramon Cleries

**Author notes:** **Corresponding author contact details**: Dr. Ville N. Pimenoff, Department of Laboratory Medicine, Karolinska Institutet, 14186, Stockholm, Sweden.

## Abstract

Viruses infecting humans are manifold and several of them provoke significant morbidity and mortality. Simulations creating large synthetic datasets from observed multiple viral strain infections in a limited population sample can be a powerful tool to infer significant pathogen occurrence and interaction patterns, particularly if limited number of observed data units is available. Here, to demonstrate diverse human papillomavirus (HPV) strain occurrence patterns, we used log-linear models combined with Bayesian framework for graphical independence network (GIN) analysis. That is, to simulate datasets based on modeling the probabilistic associations between observed viral data points, i.e different viral strain infections in a set of population samples. Our GIN analysis outperformed in precision all oversampling methods tested for simulating large synthetic viral strain-level prevalence dataset from observed set of HPVs data. Altogether, we demonstrate that network modeling is a potent tool for creating synthetic viral datasets for comprehensive pathogen occurrence and interaction pattern estimations.

## Introduction

Graphical independence network (GIN) analysis has recently been introduced as a powerful tool for identifying statistical dependencies in complex data (Djebbari and Quackenbush, 2008; Koller and Friedman, 2009; Lewis and McCormick, 2012). The most common approach to GIN is Bayesian Networks analysis (Koller and Friedman, 2009), that has been successfully applied in modeling missing data and studying probabilistic associations between multiple independent and dependent factors in human diseases (Djebbari and Quackenbush, 2008; Jansen et al., 2003; Poon et al., 2007), and also in ecological and epidemiological context (Hernán and Robins, 2006; Lewis and McCormick, 2012; Tamayo et al., 2011). In epidemiological analysis the main difference between standard multivariate regression analysis and GIN analysis is that GIN not only attempts to identify significantly associated covariates but also to disentangle if the variables are directly or indirectly dependent (Heckerman and Chickering, 1993; Koller and Friedman, 2009). Moreover, GIN analysis allows for considering each variable as conditional on the other variables, and this means that in epidemiological settings with many interrelated variables (i.e. multiple viral infections), GIN analysis might have more power to identify significant dependent patterns (e.g. co-occurrence) than standard statistical approaches (Heckerman and Chickering, 1993; Hernán and Robins, 2006; Koller and Friedman, 2009; Lewis and McCormick, 2012).

In situations, when data can be represented in a format of variables with 0/1 as presence/absence of a certain feature (variable), log-linear models can also be applied for simple assessment of relationships between variables. From this modeling technique, the most likely sub-graphs can be identified (i.e. the cliques or cluster graphs) and subsequently connecting the cliques (*i*.*e*. junction tree) can be used for simulating synthetic datasets (Cooper and Herskovits, 1992; Cowell et al., 1999; Dawid, 1992; Højsgaard, 2012; Koller and Friedman, 2009).

Currently twelve HPV infection types, i.e. HPV16/18/31/33/35/39/45/51/52/56/58/59, have been classified as carcinogenic to humans, while a group of thirteen HPV infections, i.e. HPV26/30/34/53/66/67/68/ 69/70/73/82/85/97, are considered probably or possibly carcinogenic to humans (Bouvard et al., 2009). All of these 25 HPVs are typically referred to as *high-risk* (HR) HPVs. The interaction between the above listed HPVs viral strains are not fully understood and warrants further investigation. Here, we used network modeling, based on resampling HPVs viral strain prevalence data several times and simulating a synthetic dataset from a GIN fitted to each sample set of the original data. Then each synthetic dataset is aggregated into a large database, which is subsequently used for inferring patterns of multiple associations between HPVs infections. In the study, the variables are binary (presence/absence) and consist of 42 different common HPVs viral strain infections. Our objective was to assess whether network modeling could operate as a potent tool for simulating infection networks, and thus, enabling viral strain occurrence and interaction pattern estimations at large and beyond pairwise comparison.

## Materials and methods

### Sampling and HPV prevalence

Previously published individual HPVs prevalence data from men who have sex with men (MSM, (Pimenoff et al., 2015)) and comprehensively genotyped for HPVs (covering 68 HPV strains (*ie*. types) within Alpha, Gamma, Mu and NuPVs) from anogenital wart biopsies was used as a mock of real data for this study. A total of 126 MSM with 42 different HPV type (co)-infections and known HIV status were retrieved from the published dataset (Data format shown in Table 1).

**Table 1.** Dataset modified from (Pimenoff et al., 2015) and used in our network analysis.

| ID | HIV | HPV<sub>1</sub> | HPV<sub>2</sub> | HPV<sub>3</sub> | ... | HPV<sub>41</sub> | HPV<sub>42</sub> |
| --- | --- | --- | --- | --- | --- | --- | --- |
| S001 | 0 | 0 | 0 | 1 | ... | 0 | 1 |
| S002 | 0 | 1 | 0 | 1 | ... | 0 | 0 |
| S003 | 1 | 0 | 1 | 0 | ... | 1 | 0 |
| ... | .. | .. | .. | .. | ... | .. | .. |
| S125 | 0 | 0 | 1 | 1 | ... | 0 | 1 |
| S126 | 1 | 1 | 0 | 1 | ... | 1 | 0 |

**Variable details.** **ID:** Identifier (local); **HIV:** HIV status; **HPV<sub>i</sub>:** HPV type *i* among the 42 types detected in the dataset.: **0:** No / **1:** Yes.

### Graphical modelling

In this study we used a graphical model, which is based on a directed acyclic graph (DAG), and thus, enables the estimation of a multivariate probability distribution based on conditional univariate distribution of variables (Heckerman and Chickering, 1993; Koller and Friedman, 2009). Subsequently, the DAG was converted into an undirected acyclic graph through a log-linear model (Figure 1). This was performed by first identifying a significant log-linear model and then transforming the model into an undirected graph in order to apply algorithms for deriving a probabilistic network from the graph (Figure 1) (Cooper and Herskovits, 1992; Højsgaard, 2012). Finally, these algorithms can also be used to create simulated synthetic datasets derived from the probabilistic network (Cooper and Herskovits, 1992; Højsgaard, 2012).

**Fig. 1.**
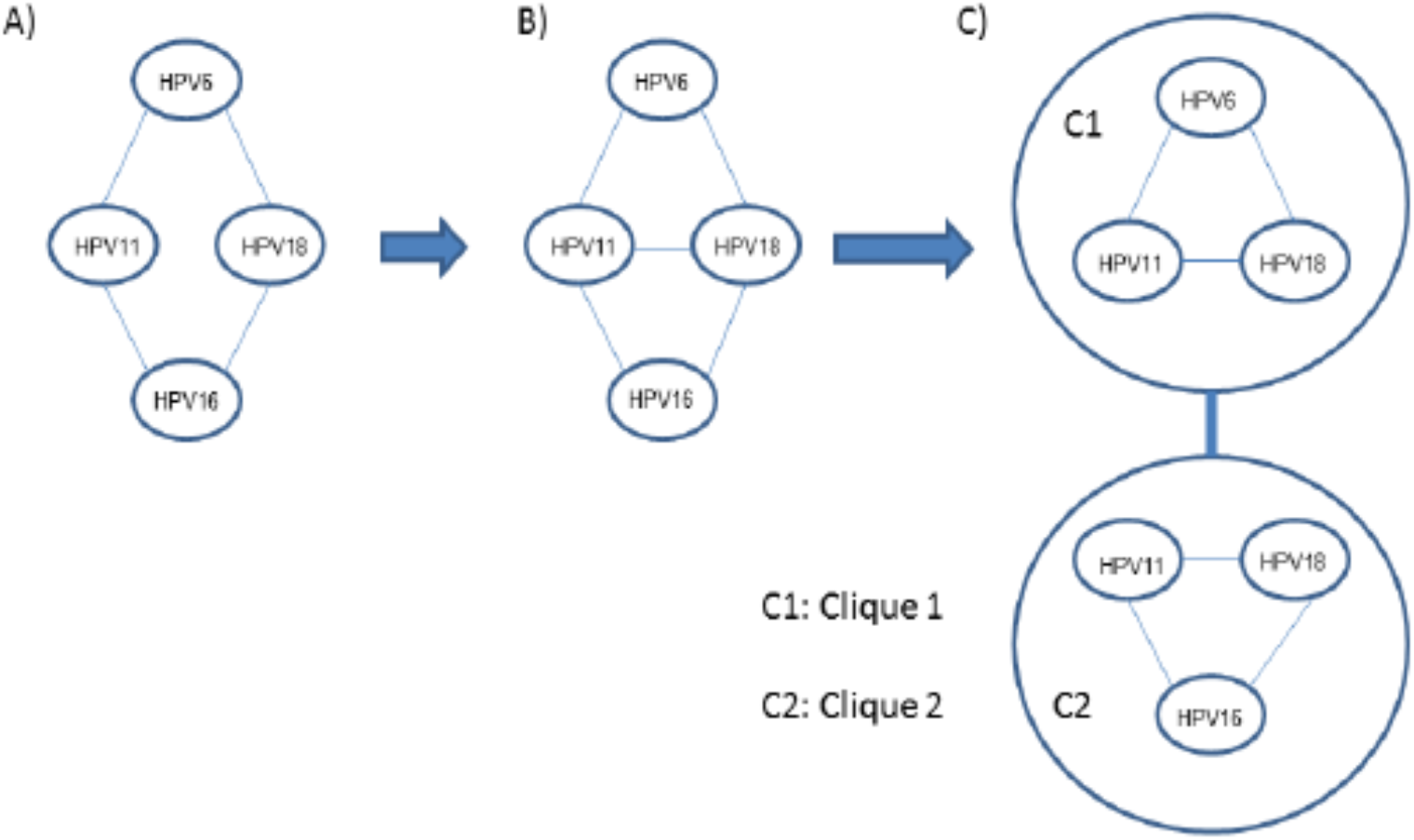
Coupling a log-linear model with a Bayesian Network, a mock example where viral strains HPV6, HPV11, HPV18 and HPV16 are considered in a log-linear model: A) Undirected graph derived from a log-linear model found by AIC; B) triangulated graph; C) junction tree and the clique potential representation of the graph. A clique tree is a junction tree if a node appears in two cliques, and it appears everywhere on the path between the cliques.

### Coupling Log-linear model with a Bayesian Network

First, we aimed to construct the association between variables through a log-linear model, as well as their independencies, and once this structure was resolved it was turned into a Bayesian network and subsequently used with junction tree algorithms to simulate a large-sample dataset (17,26). That is, *V = {HPV*_*1*_, *…, HPV*_*42*_*}* is a discrete random vector where each component is a binary variable with value 1 if HPV type is present and 0 if it is not (Table 1). Here, we assumed that we do not have missing observations across the 42 different HPV types and this vector is available for each of the *N* individuals creating a particular observed dataset (OD) matrix. Our OD is a matrix of *N* rows and 42 columns (i.e. total of different HPV types detected), which we can summarize through a contingency table (Table 1). Each cell of this table represents the count of a certain HPVs infection pattern. We can model the cell probabilities using a saturated log-linear model of the form

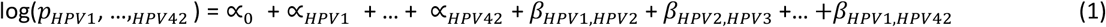

where *α* are the “main effects” and *β* the “association” terms. Suppose a mock example to introduced GIN terminology. Assume that we are interested in predicting the probability to observe an individual with the co-infection pattern described below:

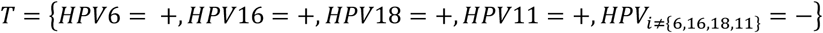

Using (1) we could estimate the probability of T, *p*_*T*_, that is

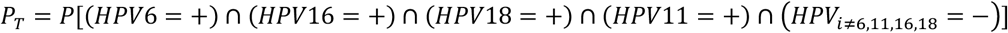

using the following equation related with a log-linear model

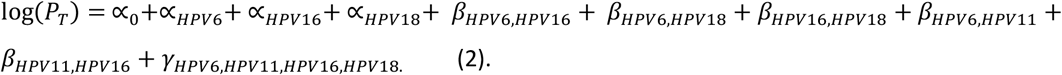

To simplify (2), *γ* parameters represent interactions of order 3 and higher. Note that terms referring to HPV types other than 6, 11, 16 and 18 are considered 0, as well as their interactions. Each variable considered in the model is a node in the graph, and each 2-variable interaction is represented by connecting 2 nodes by an arch (Fig. 1). In other words, to start with an equation (2) and using the Akaike Information Criterion (AIC) (Akaike, 1974) we can find, as an example, that the best log-linear model that fits our data is

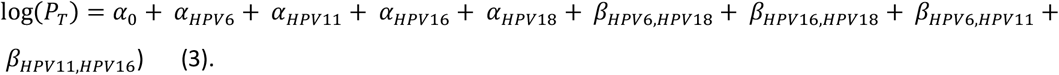

This model (3) can then be represented using an undirected graph (Fig. 1A). In GIN analysis the procedure starts with creating a triangulated graph from (3), that is coupling parent nodes and finding a graph with no cycles (Cooper and Herskovits, 1992; Højsgaard, 2012). This means that cycles of four or more nodes starting and ending at the same node cannot be found (Cooper and Herskovits, 1992; Højsgaard, 2012). Adding the arch between HPV11 and HPV18 we derive the triangulated graph (Fig. 1B). This triangulated graph is decomposable into subgraphs, or clusters of graphs (Cooper and Herskovits, 1992; Højsgaard, 2012), which are known as the “cliques” of a graph (Højsgaard, 2012) (Fig. 1C). From the cliques the key computations in GIN analysis can be performed through message passing, that is an algorithm that approximates a multivariate probability distribution by iteratively estimating marginal probability distributions of each of the variables included in the multivariate probability distribution conditional on all the other variables (Cooper and Herskovits, 1992; Højsgaard, 2012; Koller and Friedman, 2009). Then, data simulation can be performed using message passing by setting a connection between subgraphs, which is an additional graph representation, the junction tree (Fig. 1C) (Cooper and Herskovits, 1992; Cowell et al., 1999; Dawid, 1992; Højsgaard, 2012; Koller and Friedman, 2009). From the junction tree, one may obtain another representation of the joint probability distribution of interest, the clique potential representation, and based on this network structure synthetic datasets can be simulated (Cooper and Herskovits, 1992; Højsgaard, 2012).

In summary, first we searched for the best log-linear model through a backward stepwise procedure using the Akaike Information Criterion (AIC) (Akaike, 1974) and accounting for graph triangularization (no cycles) (Fig. 1A). In the second step, dependencies between variables were visualized in a graph, and then a probabilistic network was derived (Cooper and Herskovits, 1992; Højsgaard, 2012; Koller and Friedman, 2009). Once the dependency structure between HPV types was determined, a new synthetic dataset was simulated using message passing after transforming the graph network into a junction tree (Fig. 1C) (Dawid, 1992). Junction tree has a structure where nodes, variables, are branches clustering variables into cliques (17). The algorithm takes the joint probability distribution defined in the first clique and simulates values for those variables included in a given cluster according to their association, conditional probability distribution; from these values combined with the joint probability distribution of the second cluster, values for the variables are simulated in the second cluster and subsequently through the tree structure (Cowell et al., 1999; Dawid, 1992; Højsgaard, 2012).

### SynSamGIN: Procedure for generating a large-sample synthetic dataset

Since the sample size of the dataset is a limitation on deriving probabilistic patterns, we suggest a procedure for generating a large-sample synthetic dataset: **Syn**(thetic) dataset derived from **Sam**(pling) a **GIN** [SynSamGIN] fitted to the OD. This procedure, depicted in Fig. 2., has three steps: 1) To generate S databases from the OD, each one having a sample of N, the size of the original dataset, with replacement from the OD; 2) Fit a GIN model to each dataset by coupling the log-linear model selected by the AIC criterion with a BN, as described in the previous section; 3) Generate a synthetic dataset from each one of the 1..S GIN models: SynD_1_, …, SynD_S_; 4) Aggregate all SynDs into one dataset (SD).

**Fig. 2.**
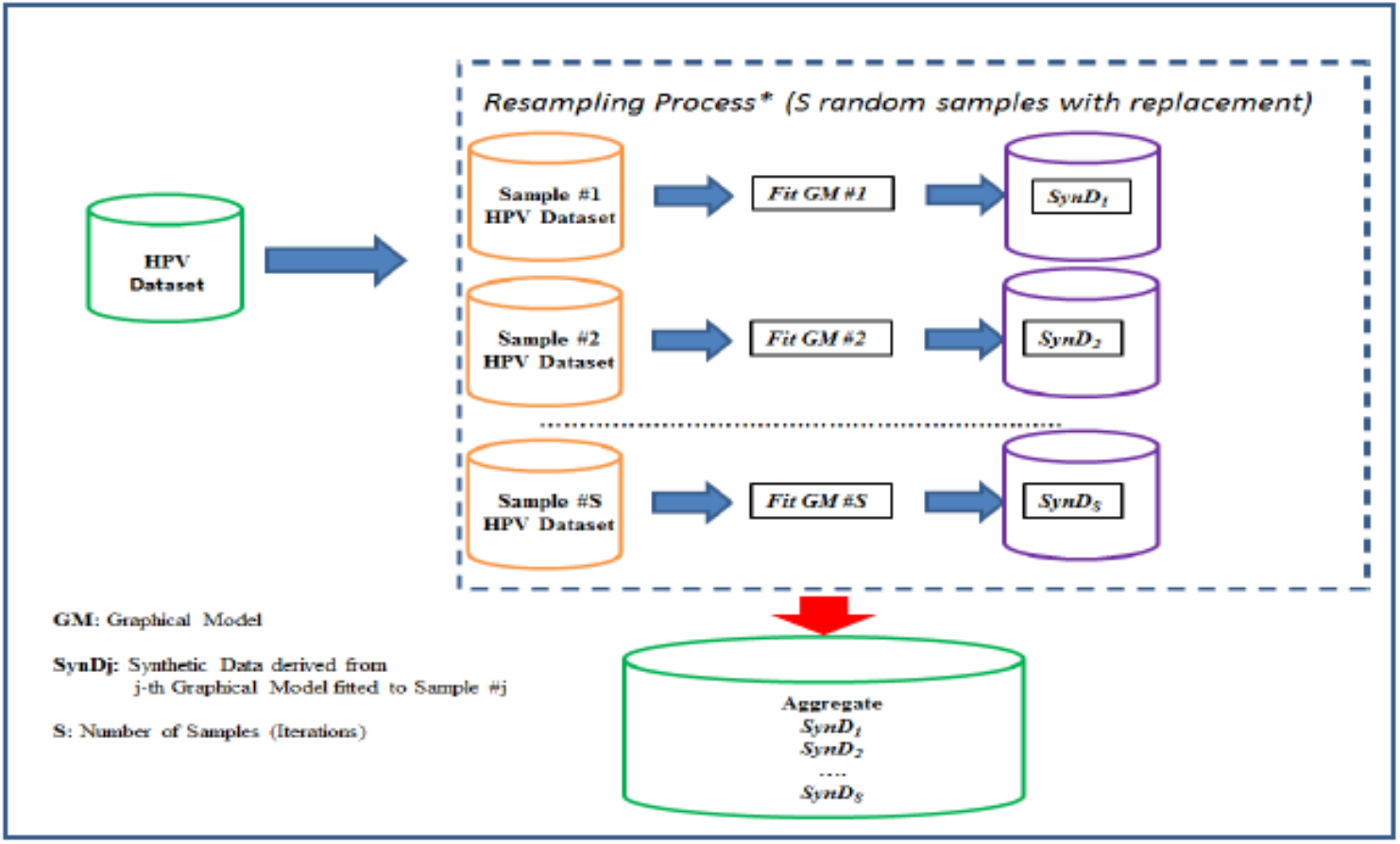
Schematic diagram for the algorithm used to generate the large sample synthetic dataset derived from the several Graphical Models fitted to resamples of the HPV Dataset.

### Simulation Study for assessing SynSamGIN procedure through a synthetic Target Population

A GIN model was fitted to an OD (N=126), and from the GIN model (Fig. 2) we generated a synthetic target population (TP) of 10,000 individuals and then, resampling from this TP we performed the simulation study. The TP included 17 HPV types, the variables, consisting of HPV6, HPV11, HPV16 and HPV18 as the highest frequency types and HPV34, HPV35 and HPV42 as the lowest type frequencies (Table 2, Full TP data as Supplementary Data S1). Hence, we compared the frequencies of each HPV-type of the TP (see Table S1) with those of obtained from generating a synthetic dataset obtained by applying *SynSamGIN* to a 500 subsets of data sampled from the TP in the three sample size scenarios. That is, for SC1 (N=50) we derived an aggregated SynD of size N=50×500=25,000, for SC2 (N=100) the SynD was N=100*500=50,000, and for SC3 (N=250) it was N=250*500=125,000 (Fig. 3). For each SynD we calculated the expected frequencies of each HPV type and derived a standardized measure for comparability reasons: the expected number of individuals out of 10,000, that is, multiplying the frequencies by 100,000 (Fig. 3). The performance of the GIN was also compared with seven commonly used oversampling algorithms: SMOTE (Synthetic Minority Over-Sampling Technique, SM) (Chawla et al., 2002) and its variant Safe-Level-SMOTE (SLS) (Bunkhumpornpat et al., 2009), Borderline-SMOTE variants 1 (BDLS1) and 2 (BDLS2), MWMOTE (Majority Weighted Minority Over-Sampling Technique, MW) (Barua et al., 2014), ADASYN (Adaptive Synthetic Sampling Approach for Imbalanced Learning) (Haibo He et al., 2008) and Random Oversampling (ROS).

**Table 2.** Frequencies and expected number of individuals (out of 10,000) in each of the seventeen HPV types considered in the TP used for the Simulation Study.

| HPV type | Freqs | Indiv. |
| --- | --- | --- |
| hvp11 | 0.3787 | 3787 |
| hvp16 | 0.1041 | 1041 |
| hvp18 | 0.0728 | 728 |
| hvp29 | 0.0100 | 100 |
| hvp31 | 0.0319 | 319 |
| hvp33 | 0.0084 | 84 |
| hvp34 | 0.0001 | 1 |
| hvp35 | 0.0001 | 1 |
| hvp39 | 0.0106 | 106 |
| hvp40 | 0.0475 | 475 |
| hvp42 | 0.0001 | 1 |
| hvp43 | 0.0391 | 391 |
| hvp44 | 0.0249 | 249 |
| hvp45 | 0.0172 | 172 |
| hvp51 | 0.0496 | 496 |
| hvp6 | 0.6533 | 6533 |
| hvp7 | 0.0410 | 410 |

Freqs: Frequencies
Indiv: Frequencies X 10,000

**Fig. 3.**
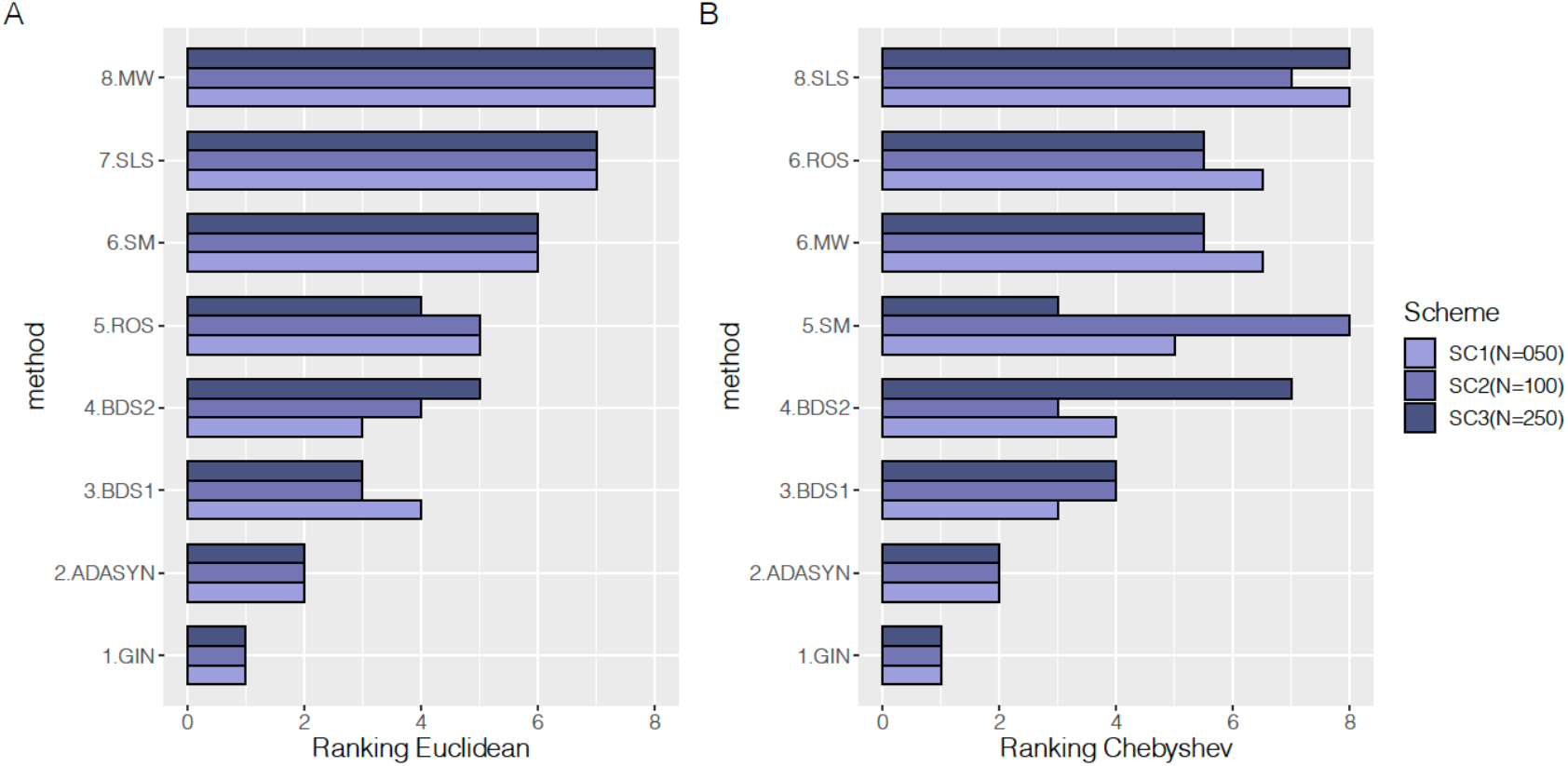
Ranking of the Euclidean (Panel A) and Chebyshev (Panel B) distances between the observed frequencies in the Target Population and the expected frequencies derived from the “synthetic” datasets generated from each method. BDSI: Borderline Smote 1; BDS2: Borderline Smote 2; MW: MWMOTE; ROS: Random Oversampling; SLS: Safe Level SMOTE; SM: SMOTE.

To assess the reliability of the synthetic datasets derived from the GINs and compared to oversampling algorithm methods, we used two metrics for the distance estimation: the Euclidean and Chebyshev distances (Cha, 2007). First 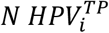 is the observed number of individuals (out of 10,000) in the TP for the i-th HPV type, whereas 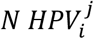 is the expected number of individuals (out of 10,000) of the i-th HPV type in the aggregated SynD derived from the j-th method (j=1,…,J=8). Second, the Euclidean distance for the j-th method is defined as 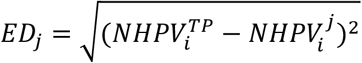, whereas the Chebyshev distance is defined as 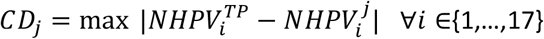. Finally, the method with smallest ED and CD is that with smallest divergence between its expected and observed frequencies in the TP.

All the analysis were carried out using R and the *gRain* R package v1.3.0 which implements the required algorithms for fitting probabilistic networks and deriving simulated datasets (Højsgaard, 2012). Hence, the package implements the core functions used by SynSamGIN described above from a triangulated undirected graph constructed from a selected log-linear model and observed dataset. Actual simulation was based on the structure of associations determined from the observed data through the triangulated graph which determined the probabilistic models (Cowell et al., 1999; Dawid, 1992) and generated the simulated dataset (SD). Here, inspecting the simulated SD included a set of new viral strain-level co-occurrence patterns predicted from the observed data. All modeling results and networks plots of inferred HPVs multiple type infections within each analysis category were carried using *gRain v1*.*3*.*0* and *gRim* v0.2.0 R packages (Højsgaard, 2012). The R code used to perform the model selection and simulations is available in GitHub PimenoffV.

## Results

To demonstrate the superior reliability of the synthetic dataset generated by GIN method, we assessed the performance of SynSamGIN under three sample size scheme SC1) N=50, SC2) N=100 and SC3) N=250 and compared the method with seven commonly used oversampling methods. Fig. 3 shows the ranking of each method in simulating a dataset from a sample of the original target population. Most importantly GIN method ranked first followed by ADASYN resampling method across all sample size scheme and methods. An additional note is that the ranking of the six remaining methods were all dependent on the sample size; for example SM method ranked third for N=250 whereas its ranking was one of the poorest for smaller sample size, if Chebyshev distances were used (Fig. 3B).

## Discussion

Our main finding is that, independently of the sample size, a synthetic viral dataset simulated using GIN and fitted to the OD outperformed any oversampling algorithms tested in this study. Hence, we present SynSamGIN, a modeling procedure for dealing with observed multiple infections prevalence data, which is used for creating a large synthetic viral prevalence datasets, which enabled the estimation of viral strain occurrence patterns not likely observed in pairwise analysis or with smaller population sample. Moreover, the advantage of using the simulated data generated from several graphical models is that one could find patterns which were not observed in the original observed data likely due to stochastic reasons. Particularly, we demonstrate the superior applicability of network modeling over resampling methods for simulating from cross-sectional viral prevalence data.

Nevertheless, the limitations of SynSamGIN is the reliability of the original data with small sample size. Moreover, a note of caution, in addition to small sample size, is that the dataset used here presents a large quantity of zeroes and this is an additional issue in finding an appropriate log-linear model to be coupled with a Bayesian network.

Taken together, our results show that a synthetic dataset simulated from a GIN fitted to the observed data could surpass oversampling algorithms for the precision of the data. We measured the divergence between the observed HPV type frequencies in a hypothetical target population versus the expected frequencies in the SynDs using Euclidean and Chebyshev divergence measures (Cha, 2007). Hence, for other type of data we also recommend to perform a similar method comparison with the corresponding divergence estimations. A recent study on fitting Bayesian network-based models has shown that computational complexity strongly depends on the class of network being learned in addition to the sparsity of the underlying DAG (Scutari et al., 2019). On the other hand, methods based on maximum likelihood estimation (MLE) of the joint distribution of the covariates from the partially classified contingency table, built using the OD, were previously assessed (Little and Rubin, 1987; Rancoita et al., 2016). However, using these methods two additional issues arose when there were huge amount of zeros in cell counts: MLE estimates were not reliable and log-linear models tended to be saturated leading to overfitting because they could not discriminate all dependencies among variables (Højsgaard, 2012; Little and Rubin, 1987; Rancoita et al., 2016; Scutari et al., 2019). Here, by the use of gRain, an offset can be added to the conditional probability tables leading to reduce zero counts and to reach smoothed probability estimates (Højsgaard, 2012). Finally, patterns derived from our simulations must be assessed and validated in future studies of larger sample size and looking for population-specific viral occurrence and interaction patterns.

Altogether estimating the true structure of the probabilistic patterns between variables (eg. infections of different viral strains) is a challenging computational problem in terms of reliable original sample size and quality of data. Our method, SynSamGIN, combines resampling and GIN analysis from the same dataset, therefore depends on the quality of the original data used. Here we have shown that the use of network modeling from a rather small but well-defined viral strain prevalence dataset was superior for data divergence comparison compared to other method using resampling. Future developments could be in a research line for implementing hybrid methodologies for preprocessing the data, and this, might decrease the computational burden and even simplify the interpretation of the probabilistic patterns related with similar viral infections’ datasets.

## Acknowledgements

The authors declare no competing interests. The authors wish to thank Jenny McCloskey for the original published HPVs prevalence data. This study was financially supported by the HEAP -study (https://heap-exposome.eu/) and RISCC -study (https://www.riscc-h2020.eu/) funded by the European Union Horizon 2020 Research and Innovation programme. None of these agencies had any role in the interpretation of the results, nor in the preparation of this manuscript.

## Notes

### Competing Interest Statement

The authors have declared no competing interest.

